## Supplementary Materials for "Microbial diversification is maintained in an experimentally evolved synthetic community"

**Supplemental materials**

**Supplemental Figures**

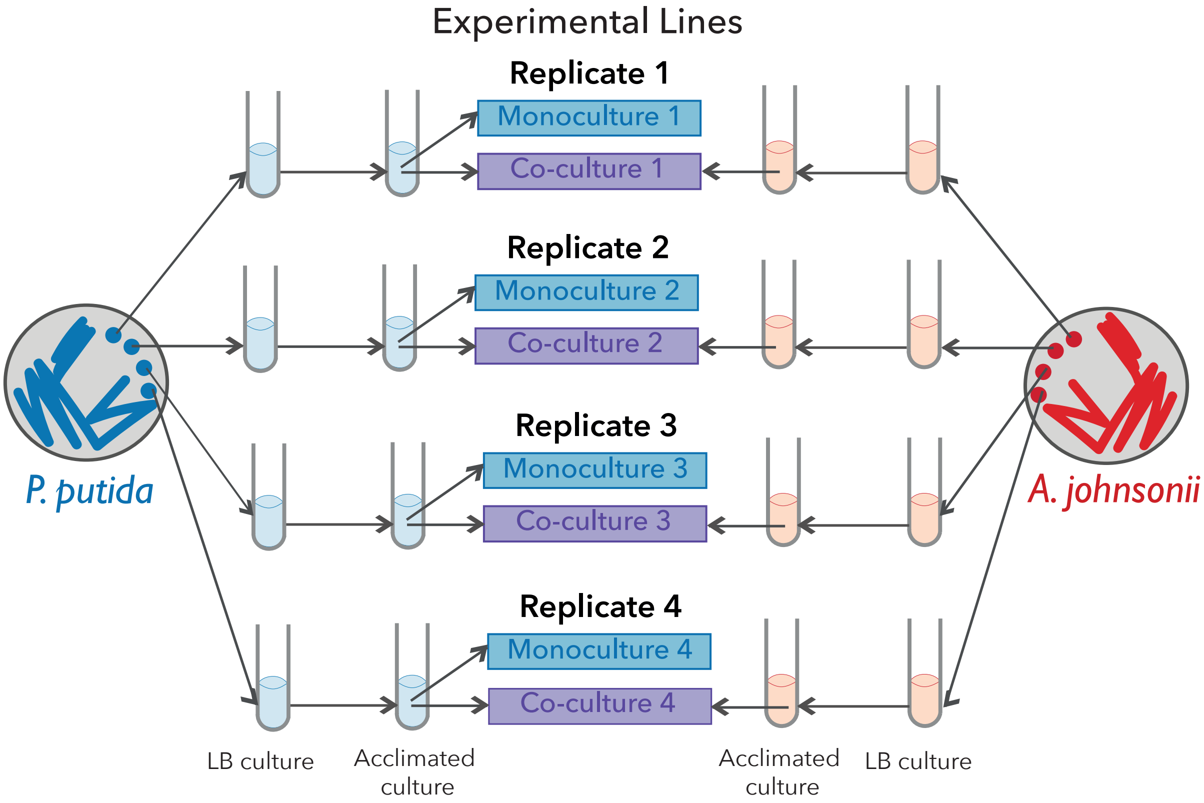


**Fig. S1. Design of the evolution experiment.** The evolution experiment started with four independent clones (i.e., four ancestors) grown in LB, washed, and acclimated in FAB medium supplemented with 0.6 mM benzoate or 0.6 mM benzyl alcohol to grow *P. putida* and A. *johnsonii*, respectively. We then mixed the acclimated cultures from each species at a ratio of 1:1 for co-cultures or evolved *P. putida* in monoculture.

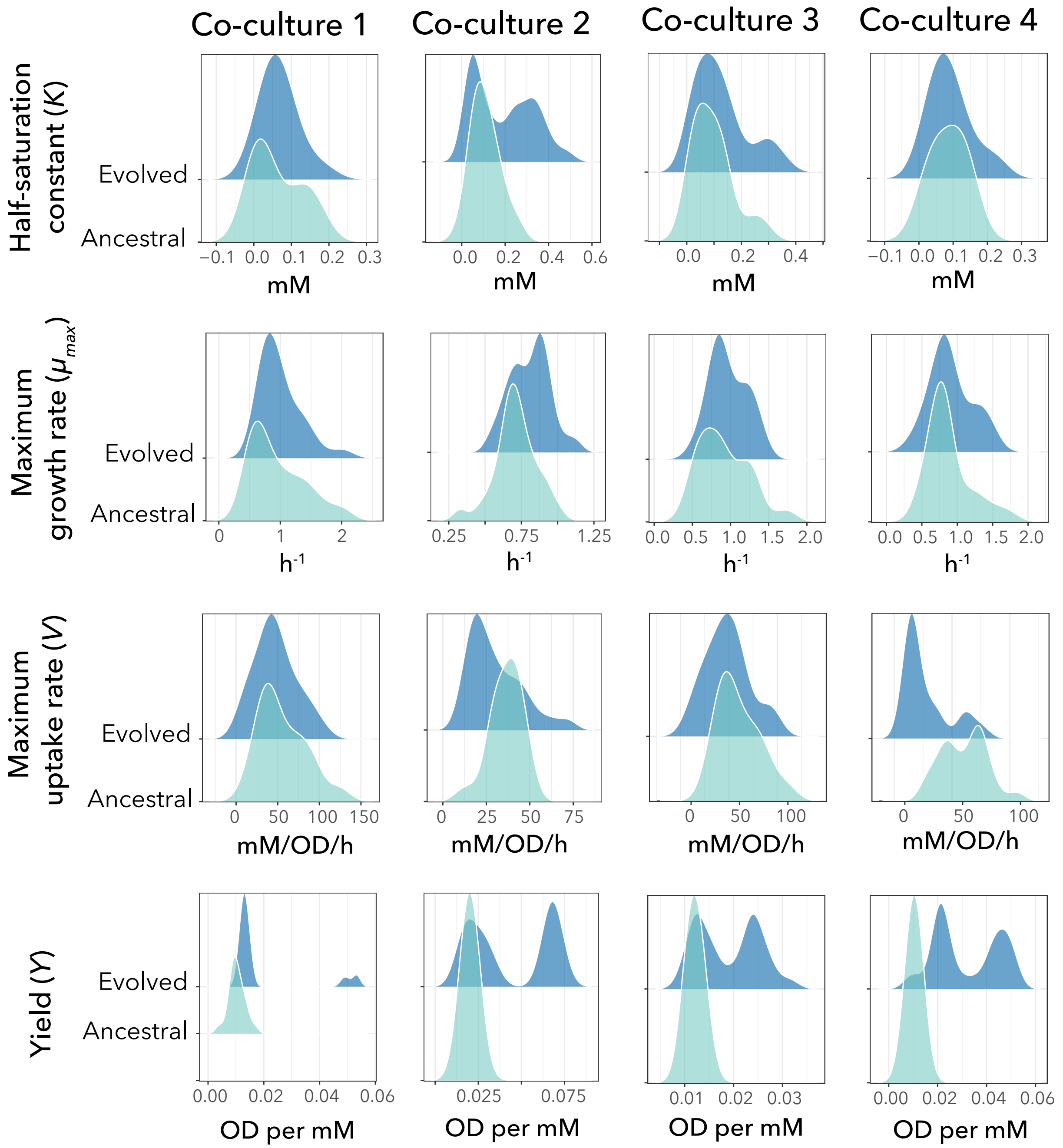


**Fig. S2. Distributions of half-saturation constant, maximum growth rate, maximum uptake rate, and yield of the ancestral and evolved *P. putida* in co-culture.** Hartigans' dip test for multimodality indicated significant bimodality in the yield of *P putida* evolved in co-culture with *p*-values equal to 0.041, 0.004, $7.48*10^{-6}$, and $2.2*10^{-16}$ for replicates 1, 2, 3, and 4 respectively.

The distribution data were plotted using the `geom_density_ridges` from the `ggridges` package in R, which smooths the data.

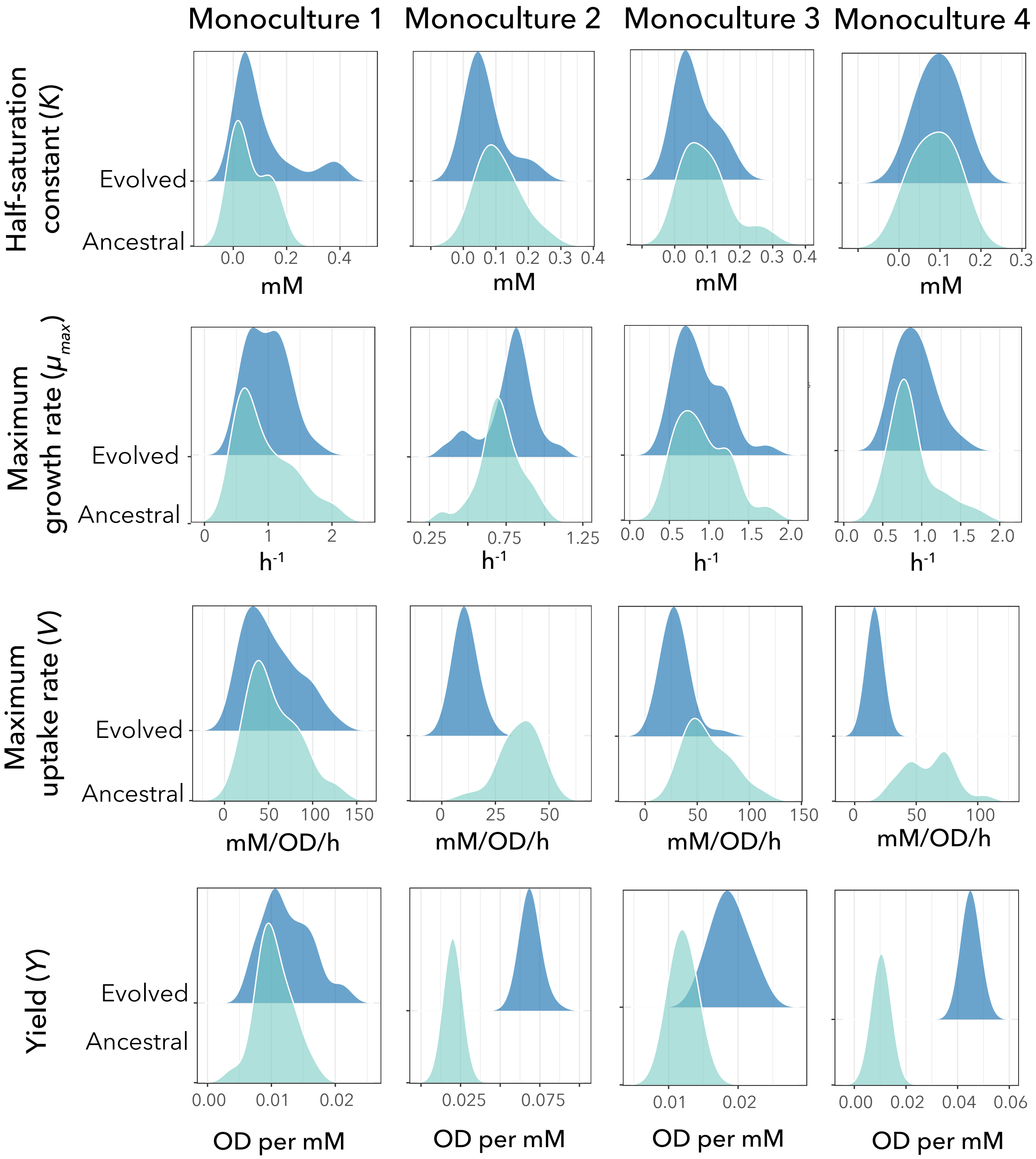


**Fig. S3. Distributions of half-saturation constant, maximum growth rate, maximum uptake rate, and yield of the ancestral and evolved *P. putida* in monoculture.** *P. putida* evolved a higher yield in monoculture compared to its ancestor based on a t-test with *p*-values equal to 0.027, $2.2*10^{-16}$, $2.2*10^{-16}$, and $2.2*10^{-16}$ for replicates 1, 2, 3, and 4, respectively. The distribution data were plotted using the `geom_density_ridges` from the `ggridges` package in R, which smooths the data.

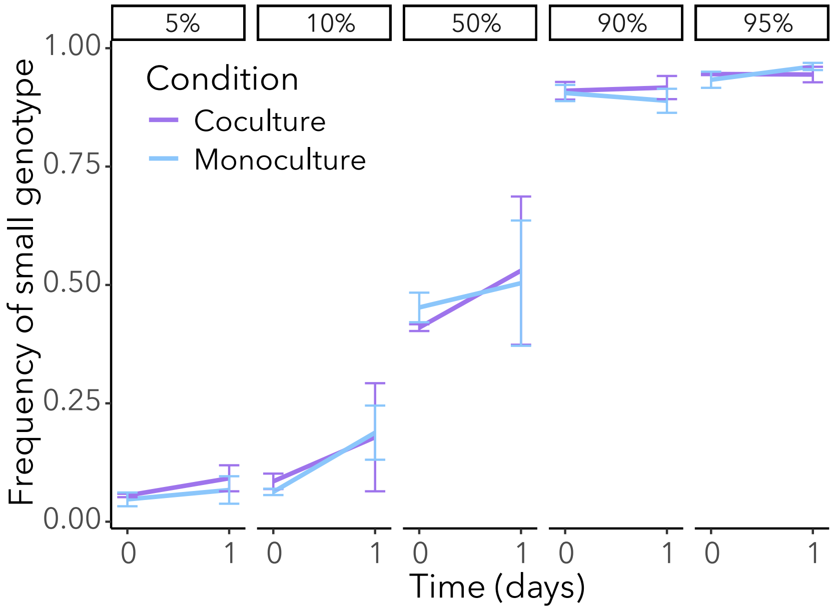


**Fig. S4. Frequency of the small genotype in head-to-head competition against the *fleQ* genotype in monoculture and co-culture over one day.** None of the genotypes can invade from rare either in monoculture or co-culture. The error bars show the standard errors of six replicate fractions (values between 0 and 1) of the small genotype.

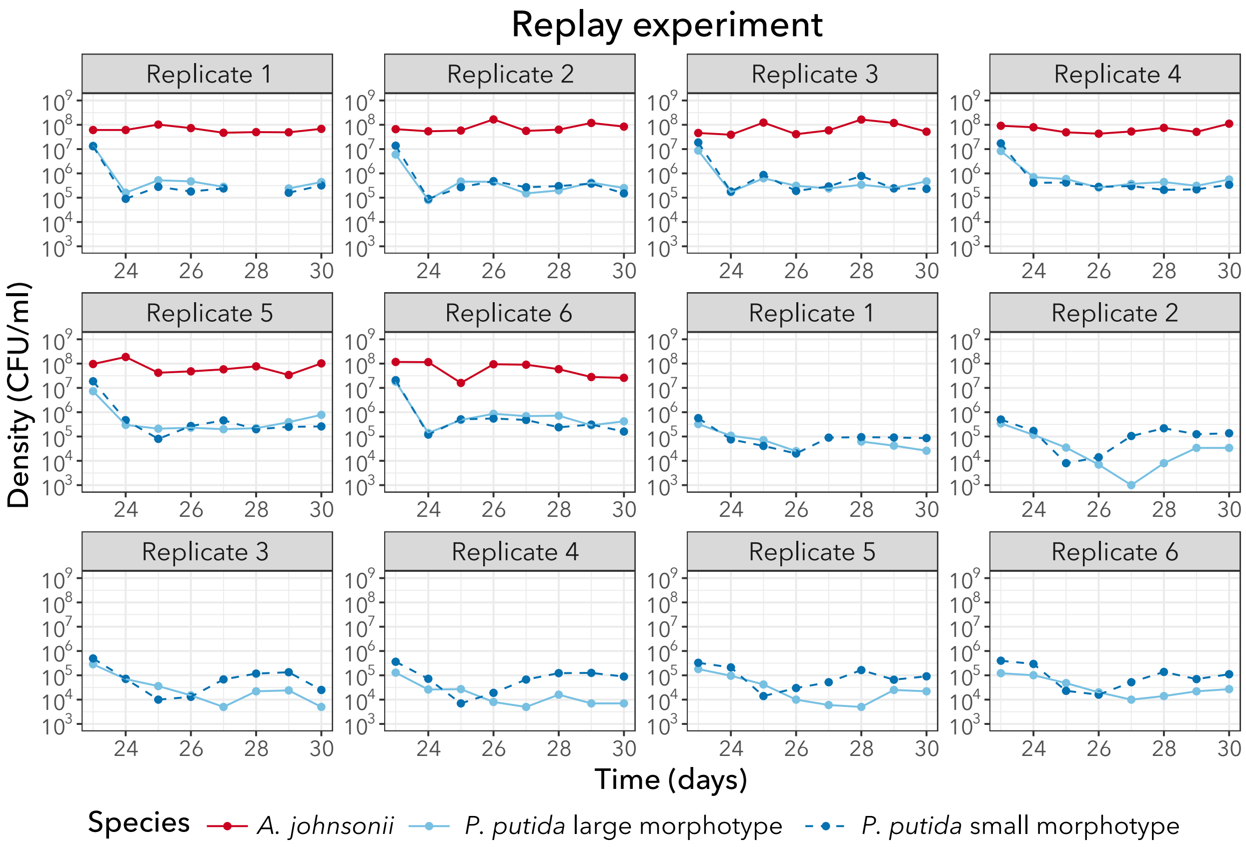


**Fig. S5. Populations trajectories of six replicates with and without *A. johnsonii* over eight days (~ 50 generations).** Each dot represents the final density (CFU mL$^{-1}$) of each type after daily cycles of growth.

### Supplementary Tables

**Table S1.** Mutations identified in populations using *breseq*.

| Position | Mutation | Freq. | gene | description |
|---|---|---|---|---|
| **Monoculture 1** | | | | |
| 1,640,523 | M143R (ATG→AGG) | 5.6% | *cmoB* ← | tRNA 5-methoxyuridine(34)/uridine 5-oxyacetic acid(34) synthase CmoB |
| 1,640,524 | M143L (ATG→CTG) | 5.2% | *cmoB* ← | tRNA 5-methoxyuridine(34)/uridine 5-oxyacetic acid(34) synthase CmoB |
| 2,768,267 | G131G (GGT→GGG) | 5.3% | *PP_RS12610* ← | TonB-dependent receptor |
| 3,282,409 | E169* (GAG→TAG) | 10.4% | *PP_RS14970* ← | cupin domain-containing protein |

| | | | | |
|---|---|---|---|---|
| 3,990,242 | +CCCGGCAG | 39.0% | *PP_RS18280* → | helix-turn-helix domain-containing protein |
| 4,647,573 | Δ130 bp | 5.5% | *[PP_RS21375]* | SulP family inorganic anion transporter |
| 6,098,739 | A117V (GCG→GTG) | 11.9% | *hexR* ← | transcriptional regulator HexR |
| **Monoculture 2** | | | | |
| 139773 | Q244P (CAG→CCG) | 5.8% | *PP_RS00675* ← | ATP-binding protein |
| 1,615,699 | A18G (GCT→GGT) | 7.3% | *PP_RS07310* ← | AbrB family transcriptional regulator |
| 4,964,741 | 2 bp→C | 100% | *fleQ* ← | transcriptional regulator FleQ |
| 5,715,528 | N65N (AAC→AAT) | 100% | *tatB* → | Sec-independent protein translocase protein TatB |
| 6,029,059 | A134A (GCC→GCA) | 100% | *PP_RS27510* ← | aldehyde dehydrogenase |
| **Monoculture 3** | | | | |
| 1,211,148 | L440R (CTG→CGG) | 5.5% | *PP_RS05530* ← | FMN-binding glutamate synthase family protein |
| 1,640,517 | I145S (ATT→AGT) | 5.5% | *cmoB* ← | tRNA 5-methoxyuridine(34)/uridine 5-oxyacetic acid(34) synthase CmoB |
| 1,640,524 | M143L (ATG→CTG) | 8.0% | *cmoB* ← | tRNA 5-methoxyuridine(34)/uridine 5-oxyacetic acid(34) synthase CmoB |
| 3,823,579 | L204R (CTG→CGG) | 5.8% | *PP_RS17595* ← | sugar kinase |
| **Monoculture 4** | | | | |
| 1,843,830 | D322H (GAC→CAC) | 100% | *PP_RS08500* ← | sensor protein GacS |
| 4,978,562:1 | +C | 100% | *flgH* ← | flagellar basal body L-ring protein FlgH |
| **Co-culture 1** | | | | |
| 1,663,184 | L403V (TTG→GTG) | 7.4% | *PP_RS07525* ← | MHS family MFS transporter |
| 1,663,192 | M400R (ATG→AGG) | 8.3% | *PP_RS07525* ← | MHS family MFS transporter |
| 3,282,468 | Δ30 bp | 26.9% | *PP_RS14970* ← | cupin domain-containing protein |
| 3,282,843 | R24H (CGC→CAC) | 6.00% | *PP_RS14970* ← | cupin domain-containing protein |
| 3,282,883 | R11G (CGT→GGT) | 15.6% | *PP_RS14970* ← | cupin domain-containing protein |
| 4,964,681 | A226V (GCC→GTC) | 23.1% | *fleQ* ← | transcriptional regulator FleQ |
| **Co-culture 2** | | | | |
| 563,729 | A271V (GCG→GTG) | 20.8% | *rpoA* → | DNA-directed RNA polymerase subunit alpha |
| 3,270,038 | F19V (TTC→GTC) | 5.4% | *PP_RS14890* ← | cupin domain-containing protein |
| 4,635,466 | G97D (GGC→GAC) | 16.9% | *uvrY* ← | UvrY/SirA/GacA family response regulator transcription factor |
| 4,964,300 | L353Q (CTG→CAG) | 35.6% | *fleQ* ← | transcriptional regulator FleQ |

| | | | | |
|---|---|---|---|---|
| **Co-culture 3** | | | | |
| 1,843,368 | G476C (GGT→TGT) | 5.1% | *PP_RS08500* ← *(gacS)* | sensor protein GacS |
| 1,844,178 | Δ1 bp | 5.2% | *PP_RS08500* ← *(gacS)* | response regulator GacS |
| 2,768,251 | F137V (TTC→GTC) | 5.3% | *PP_RS12610* ← | TonB-dependent receptor |
| 3,987,689 | Δ1 bp | 12.9% | *PP_RS18275* → *(olpA)* | hydantoinase/oxoprolinase family protein |
| 4,635,160 | T199I (ACC→ATC) | 6.3% | *uvrY* ← | UvrY/SirA/GacA family response regulator transcription factor |
| 4,635,644 | S38P (TCG→CCG) | 7.7% | *uvrY* ← | UvrY/SirA/GacA family response regulator transcription factor |
| **Co-culture 4** | | | | |
| 1,640,524 | M143L (ATG→CTG) | 5.7% | *cmoB* ← | tRNA 5-methoxyuridine(34)/uridine 5-oxyacetic acid(34) synthase CmoB |
| 4,964,290 | H356Q (CAC→CAG) | 7.5% | *fleQ* ← | transcriptional regulator FleQ |

**Table 2. Average relative growth rate of the small genotype, *i*, in head-to-head competition with the *fleQ* genotype, *j*, in monoculture or co-culture.** The values ± standard errors were obtained from three replicates.

| Starting frequency | Condition | Selection rate constant $r_{ij}$ (day$^{-1}$) | p-value* |
|---|---|---|---|
| 5% | Monoculture | 0.30 ± 0.24 | 0.328 |
| | Co-culture | 0.45 ± 0.28 | 0.251 |
| 10% | Monoculture | 1.12 ± 0.57 | 0.190 |
| | Co-culture | 0.43 ± 0.96 | 0.700 |
| 50% | Monoculture | 0.18 ± 0.60 | 0.788 |
| | Co-culture | 0.52 ± 0.73 | 0.553 |
| 90% | Monoculture | -0.17 ± 0.46 | 0.750 |
| | Co-culture | 0.16 ± 0.19 | 0.489 |
| 95% | Monoculture | 0.54 ± 0.52 | 0.402 |
| | Co-culture | 0.05 ± 0.25 | 0.848 |

* Two-sided t-test testing whether $r_{ij}$ is different than zero.
